# The senescence-inhibitory p53 isoform Δ133p53α represses the proinflammatory chemokine CXCL10 in progeria model mice and naturally aged mice

**DOI:** 10.64898/2026.03.31.715385

**Authors:** Leo Yamada, Huaitian Liu, Curtis C. Harris, Izumi Horikawa

**Affiliations:** Laboratory of Human Carcinogenesis, Center for Cancer Research, National Cancer Institute, National Institutes of Health, Bethesda, Maryland 20892, USA; CCR Collaborative Bioinformatics Resource, Center for Cancer Research, National Cancer Institute, National Institutes of Health, Bethesda, Maryland 20892, USA

**Keywords:** p53 isoform, aging, progeria mice, Luminex assay, CXCL10, proinflammatory chemokine, human GTEx dataset

## Abstract

**Aims:** Δ133p53α is a naturally occurring isoform of the human p53 protein that inhibits p53-mediated cellular senescence. We previously reported that transgenic expression of this senescence-inhibitory p53 isoform counteracts aging-associated pathological changes in progeria model mice (heterozygous *Lmna^G609G/+^*). The anti-aging effect of Δ133p53α was attributed in part to reduced levels of the proinflammatory cytokine IL-6. This study aims to comprehensively profile Δ133p53α-induced changes in cytokines and chemokines.

**Methods:** A Luminex-based multiplex quantitative assay was performed using mouse serum samples from transgenic Δ133p53α-expressing *Lmna^G609G/+^* mice and non-expressing controls. Quantitative RT-PCR and RNA *in situ* hybridization assays were used to assess Cxcl10 expression in mouse tissues. In addition, gene expression datasets from human tissues were analyzed.

**Results:** We confirmed Δ133p53α-mediated repression of serum IL-6 levels. We also found that Δ133p53α reduced serum levels of CXCL1, IL-1α, and CXCL10. We further characterized CXCL10, which has not previously been associated with progeria in mice or humans. Consistent with reduced serum CXCL10 levels, both young (15-week-old) and old (10-month-old) Δ133p53α-expressing *Lmna^G609G/+^* mice showed reduced Cxcl10 expression in the liver, spleen, and brain, major organs that produce CXCL10, compared with age-matched non-expressing controls. In naturally aged wild-type mice (2 years old), transgenic Δ133p53α expression also significantly repressed Cxcl10 expression in the spleen and brain. An inverse association between CXCL10 and Δ133p53α levels was observed in human spleen tissues, suggesting physiological relevance to human aging.

**Conclusion:** CXCL10, a proinflammatory chemokine elevated in both accelerated and natural aging, is a potential target of the anti-inflammatory activity of Δ133p53α.

## 1. Introduction

The human *TP53* gene encodes not only the full-length p53 protein (hereafter simply referred to as p53) but also multiple naturally occurring protein isoforms that are N-terminally truncated and/or C-terminally modified^[1]^. Among these, Δ133p53α is an N-terminally truncated isoform generated via alternative transcription initiated from the intron 4 promoter and alternative translation from the methionine at codon 133^[1,2]^. Δ133p53α functions as a dominant-negative inhibitory isoform of p53, preferentially inhibiting p53-mediated cellular senescence while preserving or enhancing cellular DNA repair activity^[1–4]^. We recently generated a transgenic mouse strain that enables inducible expression of this human p53 isoform to investigate its *in vivo* functions in senescence-associated diseases and conditions^[5,6]^. By crossing this strain with a progeria model (*Lmna^G609G/+^*)^[7]^, we demonstrated that Δ133p53α counteracts progeria-associated pathological changes, including those in the skin and aorta, and extends median lifespan by approximately 10%^[6]^. These anti-aging effects of Δ133p53α were primarily attributed to the widespread inhibition of cellular senescence (represented by reduced levels of p21^Waf1/Cip1^, a key effector of p53-mediated senescence, across multiple organs) and a reduction in systemic inflammation (indicated by decreased serum IL-6 levels and reduced Il6 expression in multiple organs)^[6]^.

Although IL-6 is a key cytokine contributing to systemic inflammation in both progeria-associated accelerated aging and natural aging^[8–10]^, a more comprehensive inflammatory profile requires multiplex analysis of a broad range of cytokines and chemokines. To this end, we perform a Luminex-based multiplex assay on serum samples collected from Δ133p53α-expressing *Lmna^G609G/+^* mice and non-expressing controls. This assay leads to the identification of CXCL10 as a proinflammatory chemokine that is significantly repressed by Δ133p53α. Further analyses of Cxcl10 expression in tissues from *Lmna^G609G/+^*mice and aged wild-type mice, together with interrogation of human tissue expression datasets, suggest that Δ133p53α-mediated repression of CXCL10 constitutes part of its counteracting effects on both accelerated and physiological aging.

## 2. Methods

### Mice

All animal studies have been approved by the Animal Care and Use Committee (ACUC) at National Cancer Institute (NCI, Frederick, MD) under the protocols ASP 25-264, ASP 24-466 and ASP 21-0212. All animal procedures, including housing and environmental enrichment, breeding, genotyping, recognition and alleviation of pain and distress, collection of tissues and blood, humane endpoints, and euthanasia, followed the ACUC guidelines (https://ncifrederick.cancer.gov/Lasp/ACUC/Frederick/GuidelinesFnl).

All mice used in this study were obtained in our previous study^[6]^ and their genotypes therein are given in the parentheses below. The heterozygous progeria mice included: Δ133p53α-expressing mice in which its Cre-ERT2-mediated activation was induced by tamoxifen injection at 5-6 weeks of age (*CAG-133^Tam/+^*;*Cre^Tg/+^*;*Lmna^G609G/+^*); control mice with both Δ133p53α and Cre-ERT2 transgenes but not tamoxifen-injected (*CAG-133^LSL/+^*;*Cre^Tg/+^*;*Lmna^G609G/+^*); control mice without Δ133p53α, with Cre-ERT2 and tamoxifen-injected (*CAG-133^+/+^*;*Cre^Tg/+^*;*Lmna^G609G/+^*); and control mice without Δ133p53α and Cre-ERT2 (*CAG-133^+/+^*;*Cre^+/+^*;*Lmna^G609G/+^*). The wild-type mice included: Δ133p53α-expressing mice in which its Cre-ERT2-mediated activation was induced by tamoxifen injection at 8-10 weeks of age (*CAG-133^Tam/+^*;*Cre^Tg/+^*); control mice with both Δ133p53α and Cre-ERT2 transgenes but not tamoxifen-injected (*CAG-133^LSL/+^*;*Cre^Tg/+^*); and control mice without Δ133p53α and Cre-ERT2 (*CAG-133^+/+^*;*Cre^+/+^*). All experiments used samples from 4 or 5 mice per group, with female and male mice indicated by open and closed circles, respectively, in the data presentation.

### Luminex-based multiplex quantitative assay of mouse serum samples

Serum samples were prepared from blood obtained via post-mortem cardiac puncture. Serum cytokine and chemokine levels were measured using the Mouse Cytokine/Chemokine 32-Plex Discovery Assay (MD32) (Eve Technologies, Calgary, Canada, https://www.evetechnologies.com/), a bead-based multiplex immunoassay on a Luminex platform. Assay procedures and data processing were carried out according to the manufacturer’s instructions. All samples were analyzed in duplicate to confirm the reproducibility. Statistical analyses were performed using GraphPad Prism software (version 11.0.0). *P*-values were determined by the Mann–Whitney U test.

### Quantitative RT-PCR (qRT-PCR) assays

Total RNA samples were isolated using the RNeasy Plus Micro Kit (QIAGEN, 74034). Snap-frozen tissues were homogenized in the RLT Plus buffer (supplemented with β-mercaptoethanol) provided in the kit. Cultured cells were suspended in the same RLT Plus buffer. Subsequently, both tissue and cell samples were processed according to the manufacturer’s instructions. RNA quality was checked on the Agilent TapeStation at the CCR Genomics Core, NCI. Reverse transcription for cDNA synthesis was performed using the High-Capacity cDNA Reverse Transcription Kit (Thermo Fisher Scientific, 4368814). qRT-PCR assays were performed on the 7500 Real-Time PCR system (Applied Biosystems) using the Taqman Gene Expression Master Mix (Thermo Fisher Scientific, 4369016) and the following primers/probe sets (Thermo Fisher Scientific): mouse Cxcl10 (Mm00445235_m1) and human CXCL10 (Hs00171042_m1), as well as mouse Gapdh (Mm99999915_g1) and human GAPDH (Hs02758991_g1) as normalization controls. Quantitative data analysis was performed using the ΔΔCt method (https://assets.thermofisher.com/TFS-Assets/LSG/manuals/cms_042380.pdf). All samples were analyzed in technical triplicate.

### Mouse embryonic fibroblasts (MEFs) and human fibroblasts

MEFs were prepared from mouse embryos as previously described^[6]^. Retroviral expression of progerin and 4-hydroxytamoxifen (4-OHT)-induced expression of Δ133p53α in MEFs were also performed as previously described^[6]^. Human fibroblasts from patients with Hutchinson-Gilford progeria syndrome (HGPS), AG11513 and HGADFN271, were obtained from Coriell Institute for Medical Research (https://catalog.coriell.org/) and Progeria Research Foundation (https://www.progeriaresearch.org/), respectively. Lentiviral transduction of Δ133p53α into these fibroblasts were performed as previously described^[6]^. Total RNA samples were isolated from these MEFs and human fibroblasts and analyzed by qRT-PCR as mentioned above.

### RNA *in situ* hybridization using RNAscope technology

Spleen and brain tissues were embedded in optimal cutting temperature compound (OCT) and snap-frozen. OCT-embedded tissues were cryosectioned at a thickness of 20 µm using the Cryo-Jane system (Leica Biosystems). The sections were fixed in 2% formaldehyde, and BaseScope duplex *in situ* hybridization was performed using the BaseScope™ Duplex Reagent Kit (Advanced Cell Diagnostics, #323800) according to the manufacturer’s instructions. Briefly, probes were hybridized at 40°C for 2 hours, followed by signal amplification using the provided amplification reagents. After chromogenic development, sections were counterstained with hematoxylin. The probe used to detect Δ133p53α mRNA was BA-Hs-TP53-1zz-st (Advanced Cell Diagnostics, #863951; 1 ZZ pair targeting exons 7-8 of human *TP53*; C1 channel, green), which does not cross-hybridize mouse p53 mRNA. For detecting mouse Cxcl10 mRNA, BA-Mm-Cxcl10-3EJ-C2 (Advanced Cell Diagnostics, #1560691-C2; 3 ZZ pairs targeting the exon 3 junction of mouse *Cxcl10*; C2 channel, red) was included. BaseScope images of spleen sections were scanned using the Axio Scan 2 slide scanner (Carl Zeiss Microscopy) and analyzed using QuPath v0.7.0. Cxcl10-positive staining was defined as a red chromogenic signal in the C2 channel. The percentage of Cxcl10-positive area was calculated by dividing the red-positive area by the total analyzed area, corresponding to the entire tissue section.

## Analysis of human gene expression datasets

Postmortem RNA-seq data generated by the Genotype-Tissue Expression (GTEx) consortium (v8 release) were analyzed as previously performed^[6]^. Briefly, sequencing reads were aligned to the GRCh38 human reference genome to determine transcript abundances corresponding to Δ133p53α (ENST00000504937.5, GENCODE v26) and CXCL10 (ENSG00000169245.5).

Transcripts per million (TPM) values were log -transformed [log_2_(TPM + 1)] prior to downstream analyses. Associations between Δ133p53α and CXCL10 expression levels were assessed using the Spearman rank correlation analysis. All analyses were conducted in R (version 4.5).

### Statistical analyses

Statistical analyses were performed using GraphPad Prism software (version 11.0.0). *P*-values for the serum and qRT-PCR assays were calculated using the Mann–Whitney U test or Welch’s t-test, as specified in each figure legend. For the GTEx dataset analysis, Spearman rank correlation was used to assess associations between variables. *P-*values < 0.05 were considered statistically significant.

## 3. Results

### Δ133p53α reduces serum CXCL10 levels in *Lmna^G609G/+^* progeria mice

Serum samples were prepared from 15-week-old heterozygous *Lmna^G609G/+^* progeria mice expressing transgenic Δ133p53α, along with two non-expressing control groups (no-transgene control and no-tamoxifen control) and age-matched wild-type mice (n = 4 per group). A Luminex-based multiplex quantitative assay measuring 32 mouse cytokines and chemokines was then performed (Supplementary Table S1). Consistent with our previous enzyme-linked immunosorbent assay (ELISA) data^[6]^, serum IL-6 levels were shown to be increased in *Lmna^G609G/+^*mice and decreased back to the wild-type levels by transgenic Δ133p53α expression (Figure 1A, Supplementary Table S1). Despite substantial inter-mouse variations within groups, which limited statistical significance, this multiplex assay still identified three additional factors, CXCL1 (also known as KC), IL-1α, and CXCL10 (also known as IP-10), as repressed in the Δ133p53α-expressing group, showing statistical significance relative to one control group and a borderline trend relative to the other (Figure 1B, Supplementary Table S1). This study further characterizes CXCL10, a proinflammatory chemokine associated with immune responses, chronic inflammation, and infectious diseases^[11,12]^, but which remains largely unexplored in progeria models and patients.

**Figure 1.**
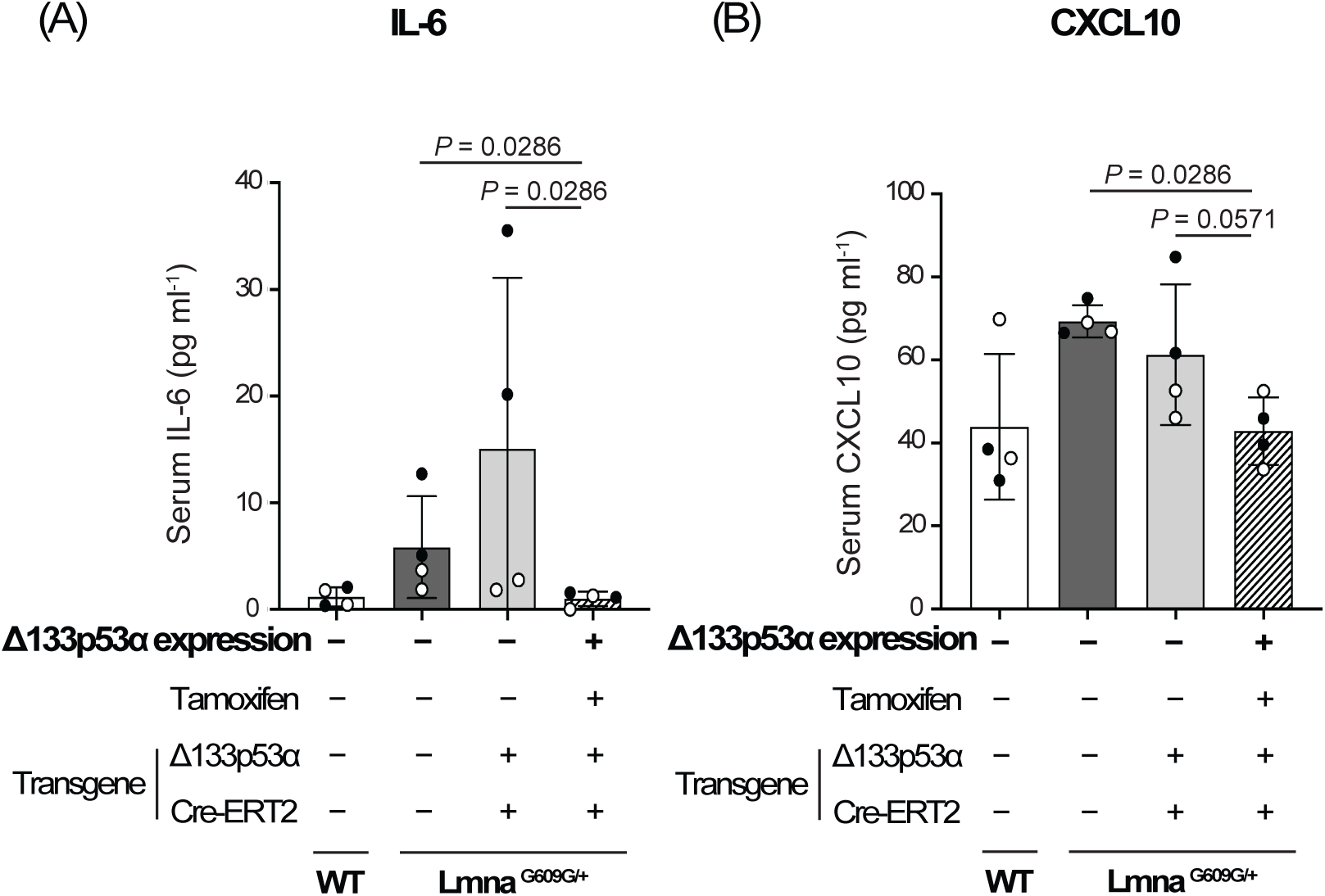
Transgenic expression of Δ133p53α reduces serum IL-6 (A) and CXCL10 (B) levels in heterozygous *Lmna^G609G/+^*progeria mice. Three groups of *Lmna^G609G/+^* mice and one group of wild-type (WT) mice at 15 weeks of age, with the indicated transgene status and tamoxifen treatment, were examined. These include a Δ133p53α-expressing *Lmna^G609G/+^* group (rightmost), two non-expressing *Lmna^G609G/+^* control groups (middle), and an age-matched WT group for comparison (leftmost). Serum concentrations (pg ml^-1^) measured using the mouse cytokine/chemokine 32-plex assay (Eve Technologies) are presented as mean ± s.d. (n = 4; open circles indicate two females, and closed circles indicate two males). *P*-values were calculated using the Mann–Whitney U test.

### Cxcl10 expression is repressed by Δ133p53α in the spleen, brain, and liver of *Lmna^G609G/+^*progeria mice

We next examined mRNA expression levels of Cxcl10 in mouse organs known to express Cxcl10, including the spleen, brain, liver, and lung^[11,12]^ (Figure 2, Supplementary Figure S1). Using the same set of 15-week-old *Lmna^G609G/+^* mice and age-matched wild-type mice as in the above serum Luminex assay (n = 4 per group), the spleen, brain, and liver showed an increase in Cxcl10 expression associated with progeria, which was reduced back to wild-type levels by transgenic Δ133p53α expression (Figures 2A-2C, left panels). These changes in Cxcl10 mRNA levels parallel the changes in serum CXCL10 levels as presented above (Figure 1B).

**Figure 2.**
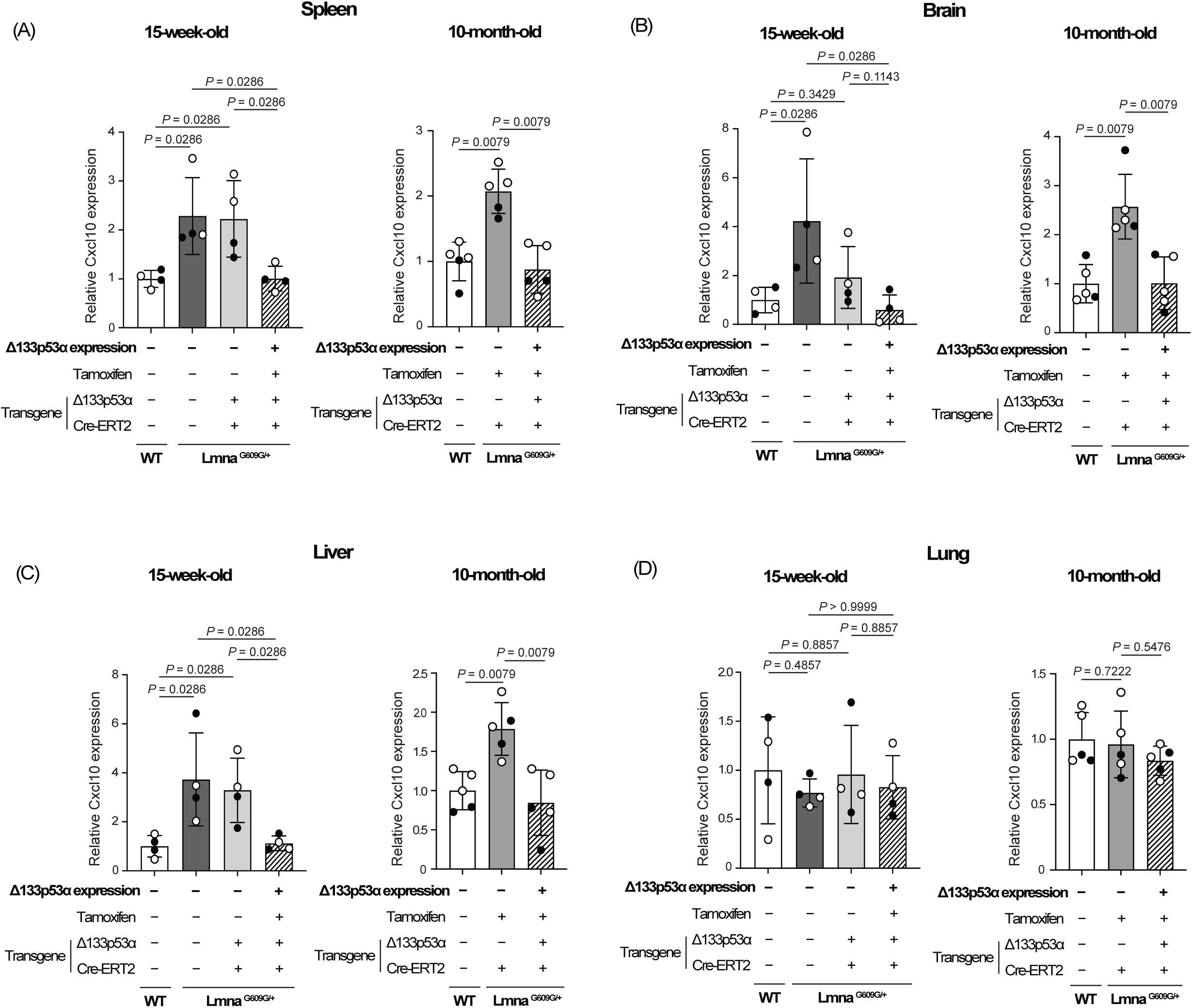
Δ133p53α represses Cxcl10 expression in the spleen, brain, and liver of *Lmna^G609G/+^*mice. Cxcl10 mRNA expression was analyzed by qRT-PCR in the spleen (A), brain (B), liver (C), and lung (D) from the same set of 15-week-old mice as in Figure 1 (left panels, n = 4), as well as from 10-month-old mice including Δ133p53α-expressing *Lmna^G609G/+^* mice, non-expressing control *Lmna^G609G/+^* mice (lacking the Δ133p53α transgene), and age-matched wild-type (WT) mice (right panels, n = 5; open circles indicate three females, and closed circles indicate two males). Data are presented as values relative to WT mice in each panel (mean ± s.d. from n = 4 or n = 5, each with technical triplicates). *P*-values were calculated using the Mann–Whitney U test.

When *Lmna^G609G/+^* mice reached 10 months of age, approaching the end of their lifespan, Cxcl10 expression levels in the spleen, brain, and liver of the Δ133p53α-expression group were again as low as those in age-matched wild-type mice, in contrast to the increased levels observed in the third non-expressing control group (tamoxifen-injected control with no Δ133p53α transgene) (Figures 2A-2C, right panels, n = 5 per group). In the lung, neither a progeria-associated increase nor a Δ133p53α-mediated decrease in Cxcl10 expression was observed at either 15 weeks or 10 months of age (Figure 2D), likely reflecting organ-specific regulations of Cxcl10 expression.

### Cxcl10 expression is repressed by Δ133p53α in the spleen and brain of naturally aged wild-type mice

To investigate the effect of Δ133p53α on Cxcl10 expression during natural aging, we examined 2-year-old wild-type mice expressing Δ133p53α, along with two age-matched control groups, no-tamoxifen control and no-transgene control (n = 4 per group). The results are presented in Figure 3, which also includes 15-week-old and 10-month-old wild-type mice (with no transgene) for comparison. Cxcl10 mRNA levels in the spleen and brain tended to increase with age, if not statistically significant (Figures 3A and 3B). In these two organs, transgenic expression of Δ133p53α significantly reduced Cxcl10 expression at 2 years of age (Figures 3A and 3B), recapitulating the results observed in *Lmna^G609G/+^* mice and suggesting that Δ133p53α regulates Cxcl10 in naturally aged mice as well. The effect of Δ133p53α on Cxcl10 expression in the liver and lung showed a weak trend but was not statistically significant (Figures 3C and 3D).

**Figure 3.**
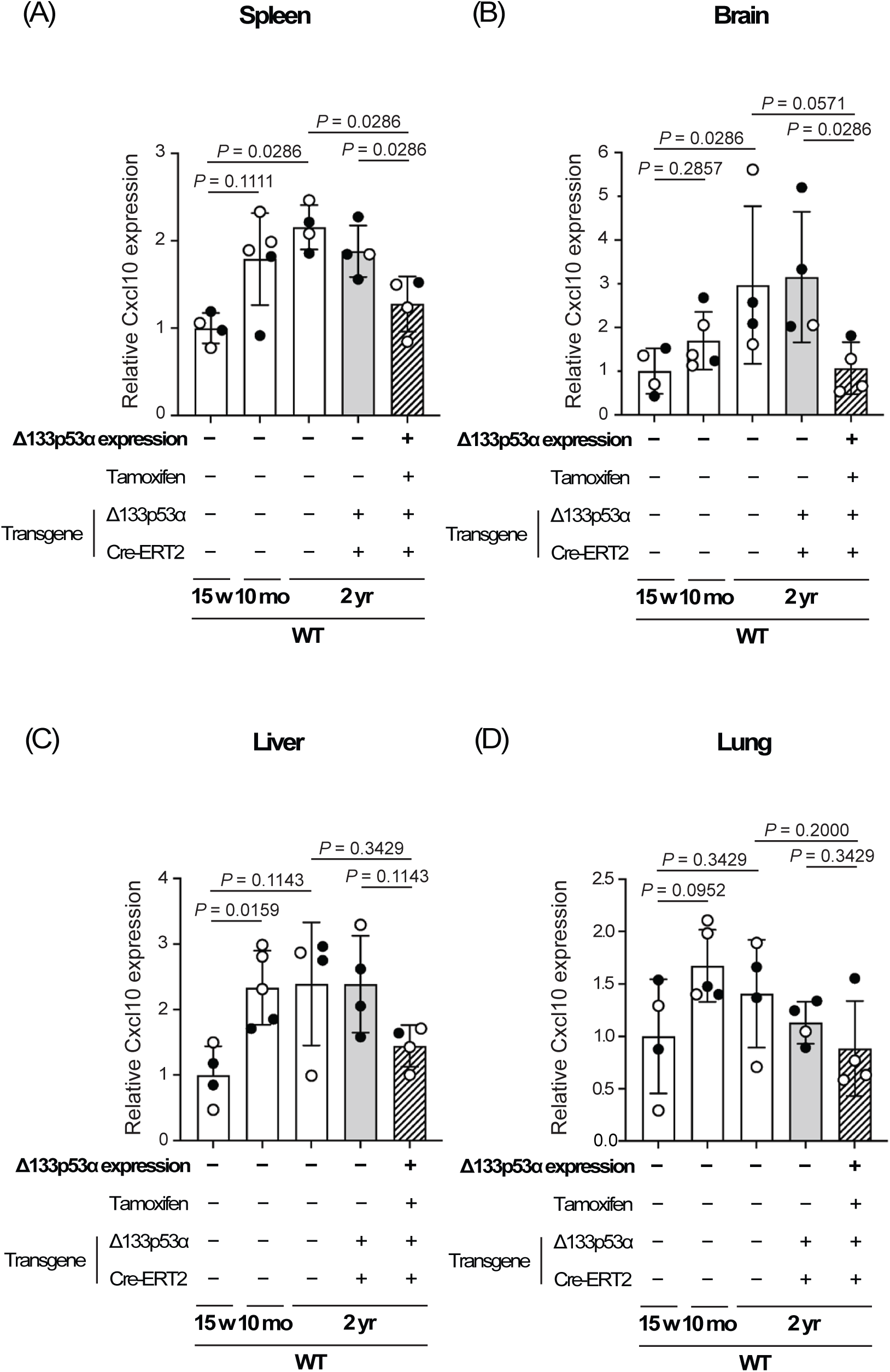
Δ133p53α represses Cxcl10 expression in the spleen and brain of naturally aged wild-type mice. Cxcl10 mRNA expression was analyzed by qRT-PCR in the spleen (A), brain (B), liver (C), and lung (D) from three groups of 2-year-old wild-type (WT) mice: a Δ133p53α- expressing group and two non-expressing control groups (no-tamoxifen and no-transgene controls) (n = 4; open circles indicate females, and closed circles indicate males). WT mice at 15 weeks and 10 months of age are shown in parallel (same data as shown above in Figure 2). All data are presented relative to 15-week-old WT mice (mean ± s.d. from n = 4 or n = 5, each with technical triplicates). *P*-values were calculated using the Mann–Whitney U test.

### Short-term expression of Δ133p53α *in vitro* represses Cxcl10/CXCL10

To perform an *in vitro* cell culture experiment with short-term expression of Δ133p53α, we established mouse embryonic fibroblasts (MEFs) from a wild-type mouse embryo carrying Δ133p53α and Cre-ERT2 transgenes. Retroviral expression of progerin significantly increased Cxcl10 mRNA levels in these MEFs (Figure 4A), recapitulating *in vivo* Cxcl10 increases observed in *Lmna^G609G/+^* mice (Figures 2A-C). Induction of Δ133p53α expression by 4-OHT for 5 days was sufficient to reverse the progerin-induced increase in Cxcl10 (Figure 4A). In two human fibroblast strains derived from HGPS patients, lentiviral expression of Δ133p53α for 5 days also significantly repressed CXCL10 mRNA levels (Figure 4B). These short-term *in vitro* effects suggest that Δ133p53α-mediated repression of Cxcl10 in mice likely occurs through direct regulation (e.g., transcriptional control), rather than through long-term *in vivo* adaptation.

**Figure 4.**
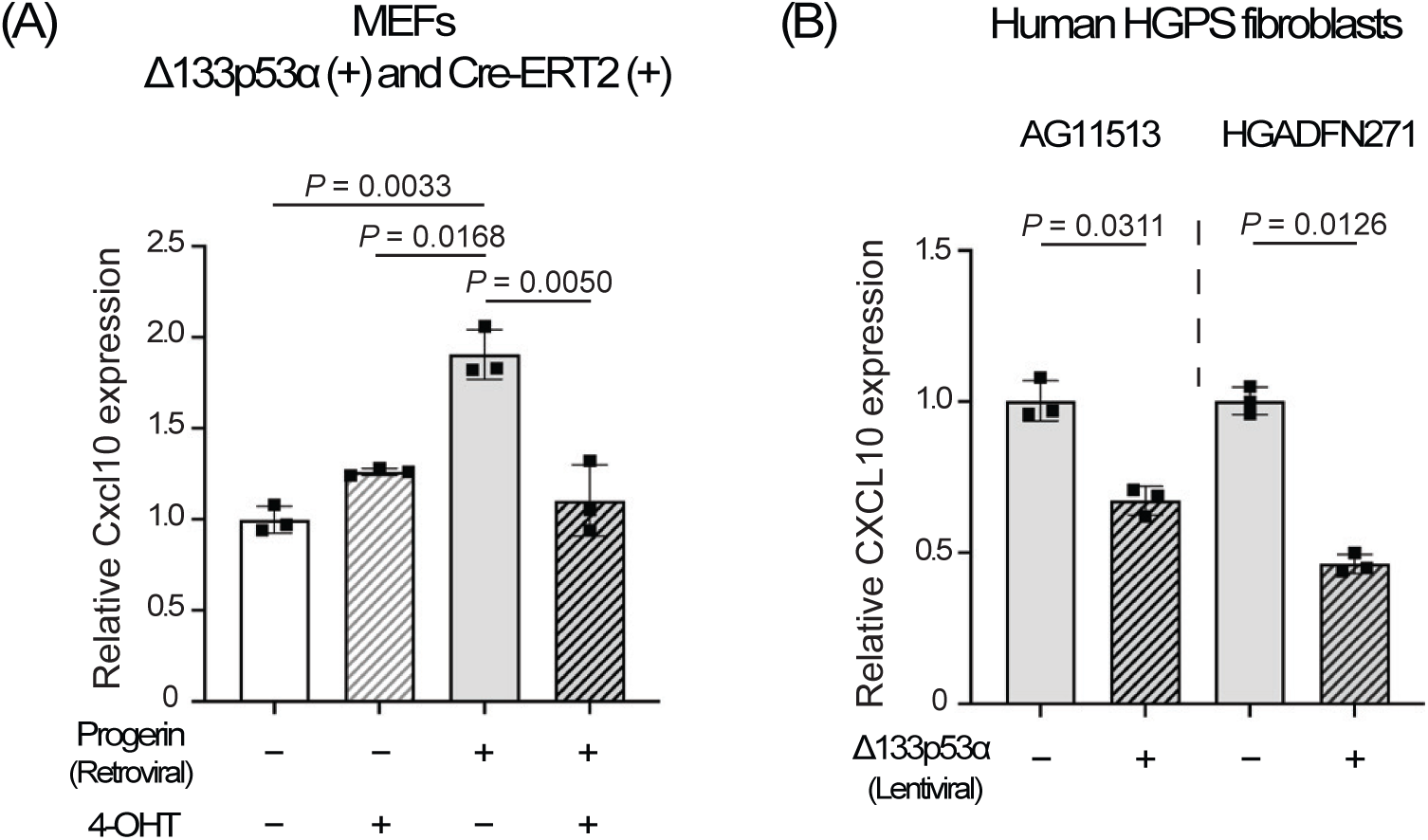
Short-term expression of Δ133p53α *in vitro* represses Cxcl10/CXCL10 expression. (A) Cxcl10 mRNA expression was analyzed by qRT-PCR in MEFs carrying both Δ133p53α and Cre-ERT2 transgenes, with or without retroviral progerin expression, and then with or without 4-OHT-induced Δ133p53α expression for 5 days. Relative expression values to progerin (-)/4-OHT (-) cells are presented as mean ± s.d. (n = 3, technical triplicates). (B) CXCL10 mRNA expression was analyzed by qRT-PCR in HGPS-derived fibroblasts, AG11513 and HGADFN271, with or without lentiviral expression of Δ133p53α for 5 days. Relative expression values to cells without Δ133p53α are presented as mean ± s.d. (n = 3, technical triplicates). *P*-values were calculated using the Welch’s t-test.

### RNA *in situ* hybridization visualizes Δ133p53α-mediated repression of Cxcl10 expression in the spleen and cerebellum

To visualize Cxcl10 and Δ133p53α expression in tissues, spleen sections from 10-month-old Δ133p53α-expressing *Lmna^G609G/+^*mice and age-matched non-expressing control mice (tamoxifen-injected, no-Δ133p53α control) were analyzed by RNAscope-based RNA *in situ* hybridization (Figure 5). As expected, positive signals for Δ133p53α mRNA (green) were observed only in Δ133p53α-expressing mice. Quantification of Cxcl10-positive signals (red) revealed significantly decreased Cxcl10 expression in Δ133p53α-expressing mice, compared with non-expressing controls (Figure 5, right). These results visually confirm the ELISA and qRT-PCR data demonstrating Δ133p53α-mediated repression of Cxcl10.

**Figure 5.**
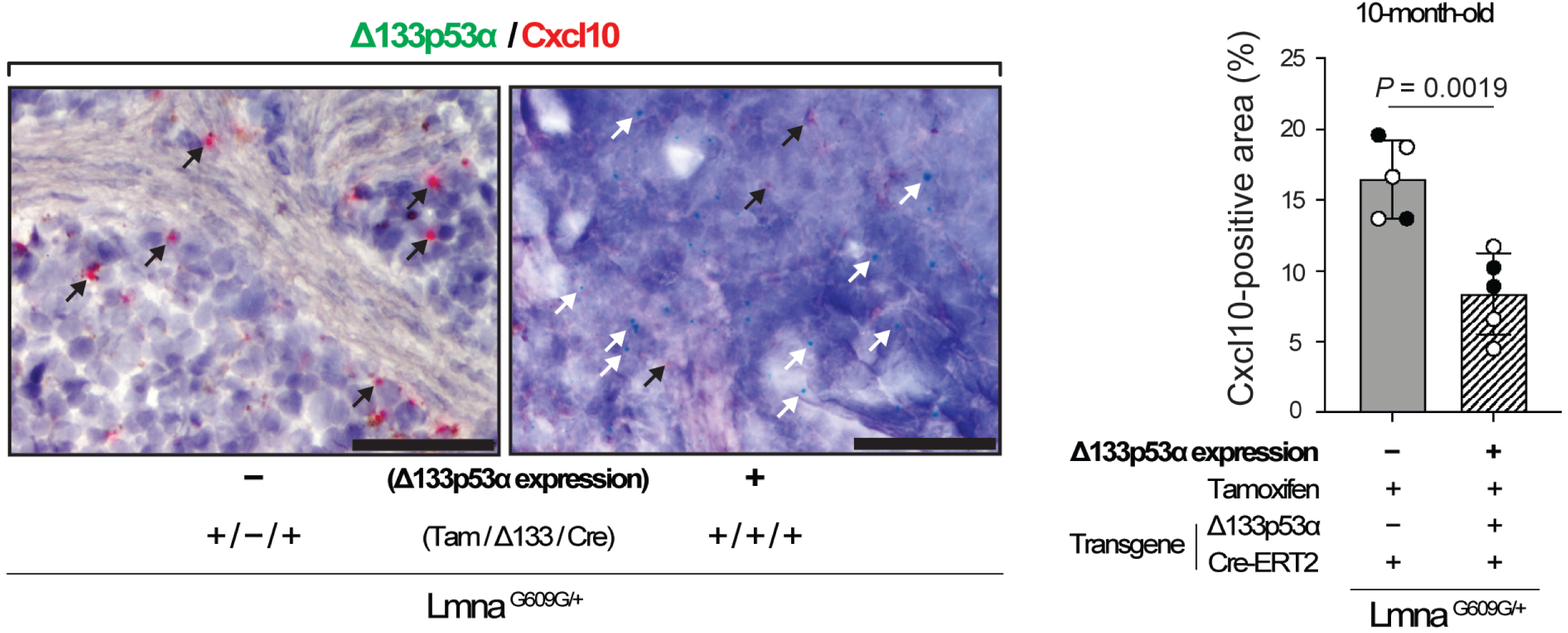
RNA *in situ* hybridization demonstrates that Δ133p53α represses Cxcl10 expression in the spleen of *Lmna^G609G/+^*mice. Spleen sections from Δ133p53α-expressing *Lmna^G609G/+^* mice (Tam/Δ133/Cre, +/+/+) and non-expressing controls (+/−/+) at 10 months of age were analyzed by BaseScope duplex *in situ* hybridization to simultaneously detect Cxcl10 mRNA (red) and Δ133p53α mRNA (green). Representative images are shown on the left. Representative Cxcl10 and Δ133p53α signals are indicated by black and white arrows, respectively. Note that Δ133p53α signals are present only in Δ133p53α-expressing mice (Tam/Δ133/Cre, +/+/+). Scale bars, 50 µm. On the right, quantitative data for Cxcl10 signals (% positive area) are presented as mean ± s.d. (n = 5; open circles indicate three females, and closed circles indicate two males). *P*-values were calculated using the Welch’s t-test.

Brain tissue sections from the same set of *Lmna^G609G/+^* mice were also examined in this assay. In non-expressing control mice, whereas the cerebral cortex showed no or only sparse staining of Cxcl10 mRNA (images not shown), the cerebellum exhibited prominent Cxcl10 staining (Supplementary Figure S2). Cxcl10-positive cell types and regions included: Purkinje neurons (Supplementary Figure S2A, left), which are known to accumulate DNA damage with age^[13,14]^; the white matter (Supplementary Figure S2B, left), where glial cells are vulnerable to aging-associated changes^[15]^; and the molecular layer (Supplementary Figure S2C, left). In Δ133p53α-expressing mice, these cell types and regions were confirmed positive for Δ133p53α mRNA, accompanied by disappearance of Cxcl10-positive cells (Supplementary Figure S2A-C, right).

### CXCL10 expression is negatively correlated with Δ133p53α expression in human spleen tissues

The repression of Cxcl10 by Δ133p53α in mice led us to hypothesize that endogenous Δ133p53α may also regulate CXCL10 expression in humans. To test this hypothesis, we examined potential associations between CXCL10 and Δ133p53α expression levels using the human GTEx RNA-seq dataset. In most organs, no significant association was observed, likely due to low expression counts and cell-type heterogeneity within the organs. Nevertheless, a significant inverse correlation between CXCL10 and Δ133p53α levels was detected in the spleen (Figure 6), which is predominantly composed of immune cells. This finding aligns with the results in mice: the spleen most reproducibly and significantly showed progeria- and aging-associated increases in Cxcl10 expression, as well as Δ133p53α-mediated repression, in *Lmna^G609G/+^*and wild-type mice (Figures 2 and 3).

**Figure 6.**
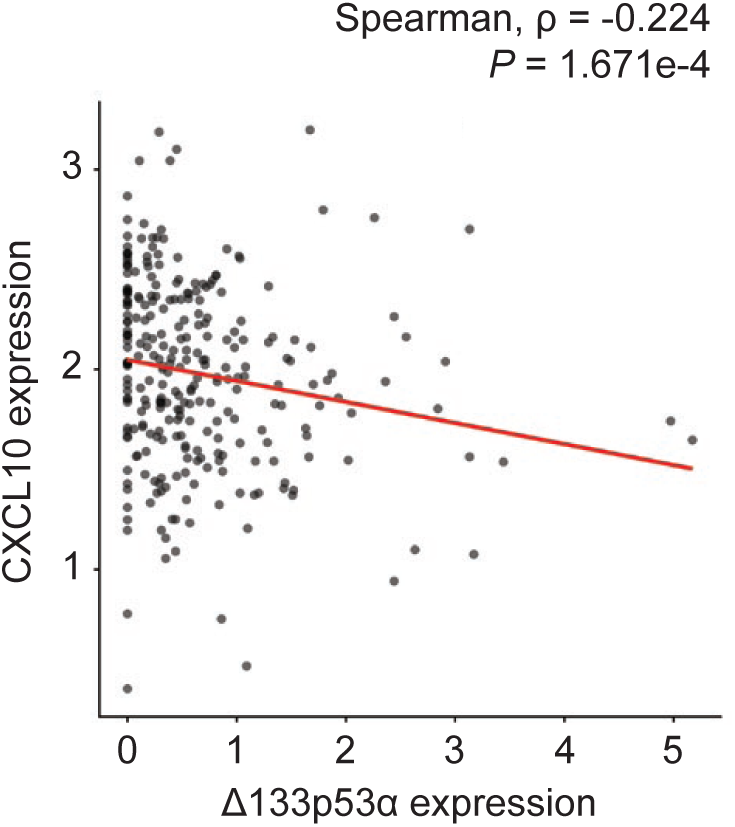
CXCL10 expression is inversely associated with Δ133p53α expression in human spleen tissues. CXCL10 and Δ133p53α expression levels in the spleen (n = 227) were obtained from analysis of the GTEx human RNA-seq dataset and are presented as log_2_(TPM + 1). Spearman’s rho and *P* value were calculated using the Spearman’s rank correlation test.

## 4. Discussion

Among various cytokines involved in the senescence-associated secretory phenotype (SASP), IL-6 plays a primary role in systemic inflammation and accelerated aging phenotypes in progeria models and patients^[6,10,16]^. A multiplex serum cytokine/chemokine assay in this study confirmed that serum IL-6 is elevated in *Lmna^G609G/+^* progeria mice and repressed by transgenic expression of Δ133p53α (Figure 1A), which counteracts accelerated aging and extends lifespan in these mice^[6]^. This multiplex assay also identified CXCL1, IL-1α, and CXCL10 as additional SASP factors repressed by Δ133p53α (Figure 1B, Supplementary Table S1). CXCL1 has been reported to mediate progeria-associated inflammation^[17]^ and to functionally interact with IL-6 through mutual activation^[18,19]^. IL-1α is known to be an upstream activator of IL-6^[20,21]^, likely enhancing progeria-associated inflammation. This study for the first time provided an opportunity to investigate CXCL10 as a progeria-linked chemokine regulated by Δ133p53α.

Consistent with a progeria-associated increase and a Δ133p53α-mediated decrease in serum CXCL10 levels, tissue expression levels of Cxcl10 in the spleen, brain, and liver were elevated in *Lmna^G609G/+^*progeria mice, compared with wild-type counterparts, and they were repressed back to wild-type levels by transgenic Δ133p53α expression (Figures 2A-2C). While CXCL10, a ligand for the CXCR3 chemokine receptor, plays a physiological role in acute infectious and inflammatory responses, its persistent production in these organs and elevated serum levels due to sustained stress, such as progeria-associated genotoxic stress, likely act in concert with increased IL-6, CXCL1, and IL-1α to promote systemic inflammation^[11,12]^. In this study, we showed that the spleen is a CXCL10-producing organ that consistently exhibits progeria- and natural aging-associated increases and Δ133p53α-mediated repression in Cxcl10/CXCL10 expression (Figures 2A, 3A, 5, 6, and Supplementary Figure S1). In addition, excessive CXCL10 production may exert deleterious local effects, which could also be mitigated by Δ133p53α. In the liver, CXCL10 and its mediated recruitment of CXCR3^+^ T lymphocytes and macrophages promote hepatic inflammation, fibrosis, and steatohepatitis^[22]^. In the brain, CXCL10 produced by reactive astrocytes and activated microglia recruits and retains CD8^+^ T lymphocytes, leading to neuroinflammation and neurodegeneration^[23,24]^. In this context, the inverse correlation between Cxcl10 and Δ133p53α observed *in situ* in the cerebellar white matter (Supplementary Figure S2B) may be consistent with our previous findings that Δ133p53α expressed in glial cells reduces SASP and neuroinflammation^[25–27]^.

Major pathological changes in progeria model mice and HGPS patients occur in the skin and the cardiovascular system including aorta^[7,28]^, where our previous study demonstrated the anti-aging effect of Δ133p53α^[6]^. In contrast, such major changes are not observed in the spleen, brain, or liver, where we showed above that Δ133p53α represses Cxcl10. We speculate that CXCL10-mediated deleterious effects in these organs do not manifest during the shortened lifespan of progeria mice and patients but may appear later in normal lifespan. In the spleen and brain of wild-type mice, we found that Cxcl10 expression tends to increase with age and is significantly repressed at 2 years of age by transgenic Δ133p53α expression (Figures 3A and 3B). These findings support the physiological and therapeutic relevance of CXCL10 and its regulation by Δ133p53α in natural aging.

The repression of Cxcl10/CXCL10 by a 5-day induction of Δ133p53α in cultured cells (Figures 4A and 4B) suggests a direct regulatory role of Δ133p53α, rather than an *in vivo* adaptation to long-term expression. Since the *CXCL10* gene has been reported to be transcriptionally activated by p53^[29]^, the dominant-negative activity of Δ133p53α against p53^[2,3]^ likely represents direct transcriptional regulation of CXCL10 by Δ133p53α, although details of the regulation remain to be elucidated. The molecular basis for the lack of progeria-associated or Δ133p53α-mediated changes in Cxcl10 expression in the lung (Figures 2D and 3D) is currently unknown, but tissue- specific differences in stress sensitivity or p53 responses to DNA damage^[30]^ may influence the Δ133p53α/p53 regulation of CXCL10.

Given that our analysis of the GTEx dataset suggests an endogenous regulation of CXCL10 by Δ133p53α in human tissues (Figure 6), this study indicates that Δ133p53α-mediated or other interventions to suppress excessive CXCL10 levels may have therapeutic potential.

Pharmacological enhancement of Δ133p53α is currently under development^[31,32]^, and a therapeutically applicable anti-CXCL10 antibody is available^[33]^. A wide range of senescence-and SASP-associated diseases could be targeted, including progeria, inflammatory liver diseases, and neurodegenerative diseases. In addition to the findings presented here, a large-scale analysis of gene expression databases^[8]^ has identified both CXCL10 and IL-6 as key biomarkers of frailty, an age-associated condition characterized by reduced physiological reserve and resilience. Combinatorial approaches targeting both CXCL10 and IL-6 are thus worth exploring.

## 5. Conclusion

CXCL10 is a proinflammatory chemokine that is elevated in both accelerated and natural aging and is repressed by the senescence-inhibitory p53 isoform Δ133p53α. The repression of CXCL10, along with IL-6, likely mediates the anti-inflammatory activity of Δ133p53α. CXCL10 may represent a therapeutic target for senescence-associated and inflammatory diseases.

## Declarations Acknowledgements

We thank Jessica Ebersole, Eleazar Vega-Valle, Morgan Whipp, Jacqueline Clem and Christine Perella for animal care and support. We also thank NCI-Frederick Molecular Histopathology Laboratory, Animal Diagnostic Laboratory, and CCR Genomics Core for their technical assistance and advice. The retroviral expression vector of progerin (pBABE-puro-GFP-progerin) was a gift from Tom Misteli through Addgene. Graphical Abstract was created in BioRender.com.

## Author contributions

Yamada L: Laboratory experiments, data analysis and interpretation.

Liu H: Data analysis and interpretation.

Harris CC: Article conception, funding acquisition.

Horikawa I: Article conception and design, data analysis and interpretation, manuscript writing.

All authors have reviewed the manuscript and approved the final version of the manuscript.

## Ethical approval

All animal studies have been approved by the Animal Care and Use Committee (ACUC) at National Cancer Institute (NCI, Frederick, MD).

## Consent to participate

Not applicable.

## Consent for publication

Not applicable.

## Availability of data and materials

The Δ133p53α-transgenic mouse strain (CAG-LSL-Δ133p53α) is maintained at the Mutant Mouse Resource and Research Center U42OD010924. The lentiviral expression vector of Δ133p53α is available at Addgene (#241916).

## Supporting information

Graphical Abstract

## References

1. Levine AJ, Hainaut P. The TP53 gene contains a diversity box that makes it more than a tumor suppressor. Cell Death Differ. 2026. DOI: 10.1038/s41418-026-01681-1

2. Fujita K, Mondal AM, Horikawa I, Nguyen GH, Kumamoto K, Sohn JJ, et al. p53 isoforms Δ133p53 and p53β are endogenous regulators of replicative cellular senescence. Nat Cell Biol. 2009;11(9):1135–1142. DOI: 10.1038/ncb1928

3. Horikawa I, Park KY, Isogaya K, Hiyoshi Y, Li H, Anami K, et al. Δ133p53 represses p53-inducible senescence genes and enhances the generation of human induced pluripotent stem cells. Cell Death Differ. 2017;24(6):1017–1028. DOI: 10.1038/cdd.2017.48

4. Roselle C, Horikawa I, Chen L, Kelly AR, Gonzales D, Da T, et al. Enhancing chimeric antigen receptor T cell therapy by modulating the p53 signaling network with Δ133p53α. Proc Natl Acad Sci U S A. 2024;121(10):e2317735121. DOI: 10.1073/pnas.2317735121

5. Nakamichi S, Yamada L, Roselle C, Horikawa I, June CH, Harris CC. The senescence-inhibitory p53 isoform Δ133p53α: enhancing cancer immunotherapy and exploring novel therapeutic approaches for senescence-associated diseases. Geroscience. 2025. DOI: 10.1007/s11357-025-01819-y

6. Yamada L, Liu H, von Muhlinen N, Harris CC, Horikawa I. Senescence-inhibitory Δ133p53α counteracts accelerated ageing and mortality. bioRxiv. 2026. DOI: 10.64898/2025.12.31.697195

7. Osorio FG, Navarro CL, Cadinanos J, Lopez-Mejia IC, Quiros PM, Bartoli C, et al. Splicing-directed therapy in a new mouse model of human accelerated aging. Sci Transl Med. 2011;3(106):106ra107. DOI: 10.1126/scitranslmed.3002847

8. Cardoso AL, Fernandes A, Aguilar-Pimentel JA, de Angelis MH, Guedes JR, Brito MA, et al. Towards frailty biomarkers: Candidates from genes and pathways regulated in aging and age-related diseases. Ageing Res Rev. 2018;47:214–277. DOI: 10.1016/j.arr.2018.07.004

9. López-Otín C, Blasco MA, Partridge L, Serrano M, Kroemer G. Hallmarks of aging: An expanding universe. Cell. 2023;186(2):243–278. DOI: 10.1016/j.cell.2022.11.001

10. Squarzoni S, Schena E, Sabatelli P, Mattioli E, Capanni C, Cenni V, et al. Interleukin-6 neutralization ameliorates symptoms in prematurely aged mice. Aging Cell. 2021;20(1):e13285. DOI: 10.1111/acel.13285

11. Groom JR, Luster AD. CXCR3 ligands: redundant, collaborative and antagonistic functions. Immunol Cell Biol. 2011;89(2):207–215. DOI: 10.1038/icb.2010.158

12. Liu M, Guo S, Hibbert JM, Jain V, Singh N, Wilson NO, et al. CXCL10/IP-10 in infectious diseases pathogenesis and potential therapeutic implications. Cytokine Growth Factor Rev. 2011;22(3):121–130. DOI: 10.1016/j.cytogfr.2011.06.001

13. Jurk D, Wang C, Miwa S, Maddick M, Korolchuk V, Tsolou A, et al. Postmitotic neurons develop a p21-dependent senescence-like phenotype driven by a DNA damage response. Aging Cell. 2012;11(6):996–1004. DOI: 10.1111/j.1474-9726.2012.00870.x

14. Shiloh Y. The cerebellar degeneration in ataxia-telangiectasia: A case for genome instability. DNA Repair (Amst). 2020;95:102950. DOI: 10.1016/j.dnarep.2020.102950

15. Hahn O, Foltz AG, Atkins M, Kedir B, Moran-Losada P, Guldner IH, et al. Atlas of the aging mouse brain reveals white matter as vulnerable foci. Cell. 2023;186(19):4117–4133.e4122. DOI: 10.1016/j.cell.2023.07.027

16. von Muhlinen N, Horikawa I, Alam F, Isogaya K, Lissa D, Vojtesek B, et al. p53 isoforms regulate premature aging in human cells. Oncogene. 2018;37(18):2379–2393. DOI: 10.1038/s41388-017-0101-3

17. Rolas L, Stein M, Barkaway A, Reglero-Real N, Sciacca E, Yaseen M, et al. Senescent endothelial cells promote pathogenic neutrophil trafficking in inflamed tissues. EMBO Rep. 2024;25(9):3842–3869. DOI: 10.1038/s44319-024-00182-x

18. Ahuja N, Andres-Hernando A, Altmann C, Bhargava R, Bacalja J, Webb RG, et al. Circulating IL-6 mediates lung injury via CXCL1 production after acute kidney injury in mice. Am J Physiol Renal Physiol. 2012;303(6):F864–872. DOI: 10.1152/ajprenal.00025.2012

19. Hou SM, Chen PC, Lin CM, Fang ML, Chi MC, Liu JF. CXCL1 contributes to IL-6 expression in osteoarthritis and rheumatoid arthritis synovial fibroblasts by CXCR2, c-Raf, MAPK, and AP-1 pathway. Arthritis Res Ther. 2020;22(1):251. DOI: 10.1186/s13075-020-02331-8

20. McCarthy DA, Clark RR, Bartling TR, Trebak M, Melendez JA. Redox control of the senescence regulator interleukin-1α and the secretory phenotype. J Biol Chem. 2013;288(45):32149–32159. DOI: 10.1074/jbc.M113.493841

21. Orjalo AV, Bhaumik D, Gengler BK, Scott GK, Campisi J. Cell surface-bound IL-1alpha is an upstream regulator of the senescence-associated IL-6/IL-8 cytokine network. Proc Natl Acad Sci U S A. 2009;106(40):17031–17036. DOI: 10.1073/pnas.0905299106

22. Xu Z, Zhang X, Lau J, Yu J. C-X-C motif chemokine 10 in non-alcoholic steatohepatitis: role as a pro-inflammatory factor and clinical implication. Expert Rev Mol Med. 2016;18:e16. DOI: 10.1017/erm.2016.16

23. Groh J, Feng R, Yuan X, Liu L, Klein D, Hutahaean G, et al. Microglia activation orchestrates CXCL10-mediated CD8(+) T cell recruitment to promote aging-related white matter degeneration. Nat Neurosci. 2025;28(6):1160–1173. DOI: 10.1038/s41593-025-01955-w

24. Lawrence JM, Schardien K, Wigdahl B, Nonnemacher MR. Roles of neuropathology-associated reactive astrocytes: a systematic review. Acta Neuropathol Commun. 2023;11(1):42. DOI: 10.1186/s40478-023-01526-9

25. Turnquist C, Beck JA, Horikawa I, Obiorah IE, von Muhlinen N, Vojtesek B, et al. Radiation-induced astrocyte senescence is rescued by Δ133p53. Neuro Oncol. 2019;21(4):474–485. DOI: 10.1093/neuonc/noz001

26. Turnquist C, Horikawa I, Foran E, Major EO, Vojtesek B, Lane DP, et al. p53 isoforms regulate astrocyte-mediated neuroprotection and neurodegeneration. Cell Death Differ. 2016;23(9):1515–1528. DOI: 10.1038/cdd.2016.37

27. Ungerleider K, Beck JA, Lissa D, Joruiz S, Horikawa I, Harris CC. Δ133p53α protects human astrocytes from amyloid-beta induced senescence and neurotoxicity. Neuroscience. 2022;498:190–202. DOI: 10.1016/j.neuroscience.2022.06.004

28. Hennekam RC. Hutchinson-Gilford progeria syndrome: review of the phenotype. Am J Med Genet A. 2006;140(23):2603–2624. DOI: 10.1002/ajmg.a.31346

29. Wahyuni T, Tanaka S, Igarashi R, Miyake Y, Yamamoto A, Mori S, et al. CXCL10 is a novel anti-angiogenic factor downstream of p53 in cardiomyocytes. Physiol Rep. 2022;10(9):e15304. DOI: 10.14814/phy2.15304

30. Hill RJ, Bona N, Smink J, Webb HK, Crisp A, Garaycoechea JI, et al. p53 regulates diverse tissue-specific outcomes to endogenous DNA damage in mice. Nat Commun. 2024;15(1):2518. DOI: 10.1038/s41467-024-46844-1

31. Joruiz SM, Lissa D, von Muhlinen N, Dranchak PK, Inglese J, Horikawa I, et al. Pharmacologic activation of Δ133p53α reduces cellular senescence in progeria patients-derived cells. Aging Pathobiol Ther. 2025;7(3):231–238. DOI: 10.31491/APT.2025.09.185

32. Lissa D, Joruiz SM, Dranchak PK, Ungerleider K, Yamada L, Horikawa I, et al. A quantitative high-throughput screen identifies compounds that upregulate the p53 isoform Δ133p53α and inhibit cellular senescence. ACS Pharmacol Transl Sci. 2025;8:2061–2074. DOI: 10.1021/acsptsci.5c00186

33. Sandborn WJ, Colombel JF, Ghosh S, Sands BE, Dryden G, Hébuterne X, et al. Eldelumab [Anti-IP-10] induction therapy for ulcerative colitis: a randomised, placebo-controlled, phase 2b study. J Crohns Colitis. 2016;10(4):418–428. DOI: 10.1093/ecco-jcc/jjv224

