## Supplementary figures and images for "The senescence-inhibitory p53 isoform Δ133p53α represses the proinflammatory chemokine CXCL10 in progeria model mice and naturally aged mice"

### Graphical Abstract

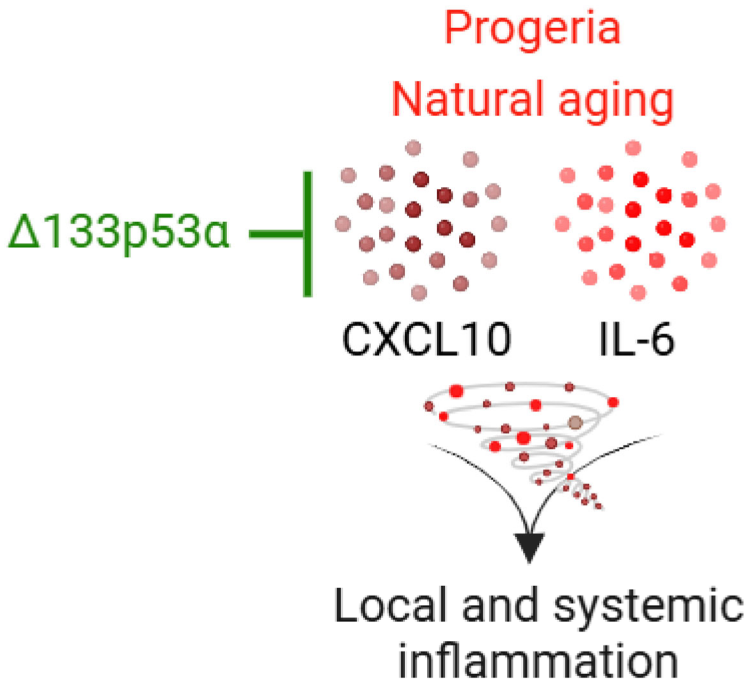
