## Supplementary Figures S1 and S2 for "The senescence-inhibitory p53 isoform Δ133p53α represses the proinflammatory chemokine CXCL10 in progeria model mice and naturally aged mice"

**Supplementary Figure S1**

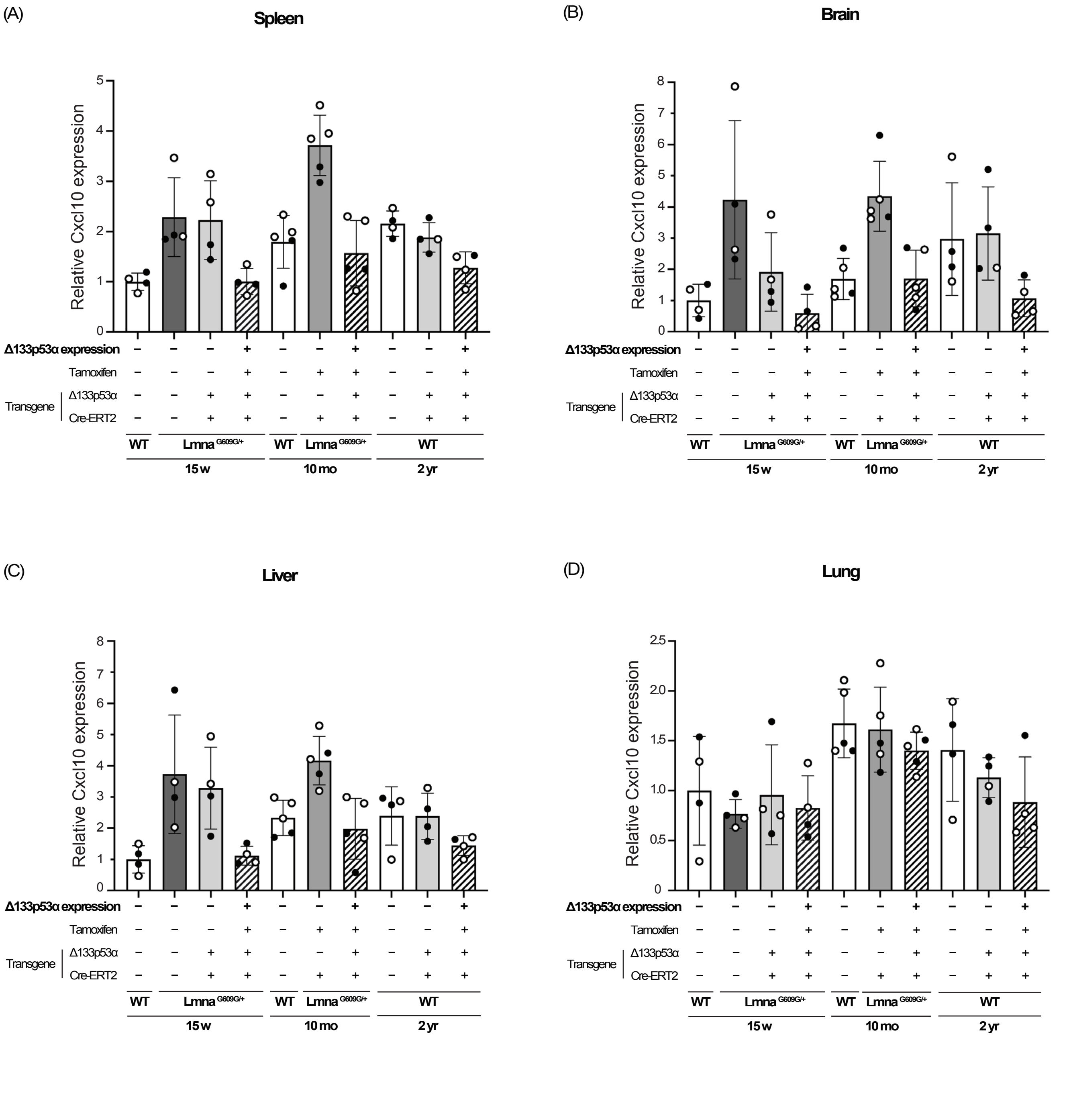

**Supplementary Figure S1.** All qRT-PCR data in the spleen (A), brain (B), liver (C), and lung (D). Data from *Lmna*$^{G609G/+}$ and wild-type (WT) mice at different ages are shown together for cross-reference. Transgene status and tamoxifen treatment are indicated as in Figures 2 and 3. All data (mean ± s.d.) are presented relative to 15-week-old WT mice (n = 4 or 5; open circles indicate female mice, and closed circles indicate male mice). *P* values are shown in each panel of Figures 2 and 3.

Supplementary Figure S2

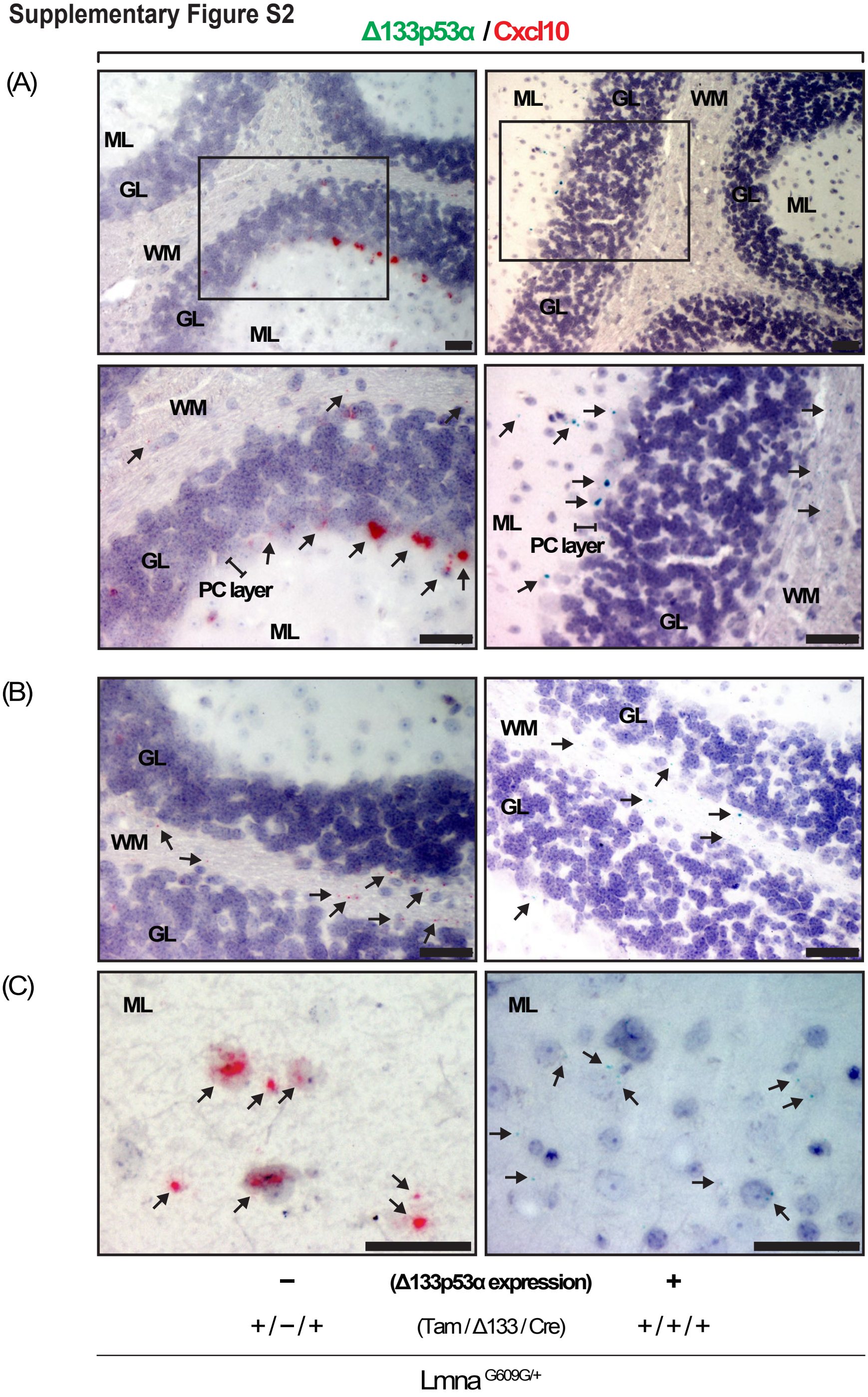

**Supplementary Figure S2.** RNA *in situ* hybridization shows that Δ133p53α represses Cxcl10 expression in the cerebellum of *Lmna*$^{G609G/+}$ mice. Brain sections from Δ133p53α-expressing *Lmna*$^{G609G/+}$ mice (Tam/Δ133/Cre, +/+/+) and non-expressing controls (+/–/+) at 10 months of age were analyzed by BaseScope duplex *in situ* hybridization to simultaneously detect Δ133p53α mRNA (green) and Cxcl10 mRNA (red). Presented are images of the cerebellum containing the Purkinje cell (PC) layer (A; insets are shown enlarged), the white matter (WM) (B), and the molecular layer (ML) (C). GL, granular layer. Magnification, 10× (A, upper), 20× (A, lower, and B), and 40× (C). Scale bars, 50 µm. Arrows indicate positive signals, appearing as small punctate dots or dense clusters. In (B) and (C), the exact identities of positively stained cell types are not clearly defined, although they are likely oligodendrocytes, astrocytes, microglia, or oligodendrocyte precursor cells in (B), and basket cells or stellate cells in (C).
